# CnidoSite: A Multi-Omics Database for Cnidarians

**DOI:** 10.64898/2026.10.15.758670

**Authors:** Jiajie She, Changsen Jiang, Jiakai Fan, Pei-Yuan Qian, Longjun Wu

## Abstract

The phylum Cnidaria, encompassing reef-building corals, sea anemones, hydroids, and jellyfish, includes both ecological keystones of the marine biosphere and indispensable models for reconstructing early animal evolution. However, escalating ocean warming, recurrent mass coral bleaching events, and intensifying jellyfish blooms have underscored the urgent need to dissect the molecular underpinnings of cnidarian thermal tolerance, symbiosis, and environmental adaptation. Progress in cnidarian research is hindered by the lack of centralized, high-quality multi-omics resources. To address this, we present CnidoSite (https://CnidoSite.org), a comprehensive, analysis-ready multi-omics platform dedicated to Cnidaria. Spanning 327 species across 120 families, 23 orders, and 7 classes, it integrates 147 full-annotated genomes, 15,642 bulk transcriptomes, 50 single-cell datasets, 46 proteomes, 723 epigenomes, 1,650 metagenomic samples, and 180,652 phenotype records. Complemented by advanced toolkits for multidimensional functional and comparative analyses, CnidoSite provides extensive functional genomic resources, including genome assemblies, phylogenies, fossil entries, functional annotations, BUSCO assessments, and mitochondrial annotations. Key features include 29 co-expression networks with cross-tissue visualization and single-cell atlases to map developmental trajectories and stress responses. In summary, CnidoSite serves as a centralized platform that bridges cnidarian biology, marine conservation, and evolutionary genomics to investigate coral bleaching, symbiosis breakdown, jellyfish blooms, and the mechanisms driving environmental adaptation.

## Introduction

The phylum Cnidaria—comprising reef-building corals, jellyfish, sea anemones, and hydroids—ranks among the most ecologically pivotal and evolutionarily significant animal lineages. Ecologically, cnidarians form the backbone of global marine ecosystems: scleractinian corals sustain roughly 25% of marine biodiversity and support the livelihoods of nearly one billion people (1), while medusa-forming cnidarians (jellyfish and hydroids) occupy critical nodes in pelagic food webs. Both groups now face unprecedented climate-driven stress. Recurrent mass bleaching has triggered widespread tropical reef degradation and fundamentally restructured benthic assemblages (2–5), whereas intensifying jellyfish blooms are disrupting pelagic ecosystems and imposing substantial economic costs on fisheries, tourism, and coastal infrastructure (6). These converging threats underscore the urgent need to elucidate the molecular mechanisms governing cnidarian thermal tolerance, host–symbiont recognition, and environmental adaptation. From an evolutionary perspective, as one of the earliest-branching metazoan lineages, Cnidaria provides an indispensable model for deciphering the origins and evolution of multicellularity, tissue organization, and developmental complexity (7–10).

Advances in high-throughput sequencing have catalyzed substantial progress in cnidarian genomics. Since the first cnidarian genome was published (*Nematostella vectensis*) (11), over 300 cnidarian genomes have been released, vastly expanding the genomic landscape of this phylum. These resources have since been enriched by transcriptomic, epigenomic, proteomic, and metagenomic datasets, alongside functional studies identifying candidate genes underlying thermal resilience and venom production (12, 13). Single-cell transcriptomic atlases have since resolved cellular heterogeneity of cnidarian tissues, mapping distinct cell states during development, regeneration, and stress responses (14, 15). Despite this data explosion, integration remains a critical bottleneck. While early efforts such as CnidBase provided foundational, gene-centric resources for comparative evolutionary analyses (16), multi-omics data remain scattered across disparate repositories—including NCBI, Ensembl, the National Genomics Data Center, and the Reefgenomics database—each employing inconsistent annotation standards, data formats, and quality-control thresholds (17–19). This fragmentation severely impedes data discovery, cross-study comparison, and systematic integrative analysis, posing a major barrier to the cnidarian research community.

To fill this critical gap, we developed CnidoSite, a comprehensive, analysis-ready multi-omics platform purpose-built for the phylum Cnidaria. Currently, CnidoSite curates data across 327 species spanning 120 families, 23 orders, and 7 classes. This platform integrates 147 full-annotated genome, 15,642 bulk transcriptomes, 50 single-cell transcriptomic datasets, 46 proteomes, 723 epigenomes, 1,650 metagenomic samples, and 180,652 phenotype records, including bleaching traits and thermal tolerance metrics. Beyond serving as a multi-omics data repository, CnidoSite adds functional-genomic layers to these resources, including phylogenies, fossil records, BUSCO evaluations, and mitochondrial annotations. Key features of CnidoSite include 29 co-expression networks and a fine-grained single-cell atlas annotated with curated cell markers, enabling detailed mapping of developmental trajectories and stress-responsive landscapes. By integrating these extensive datasets with multi-dimensional analytical tools, CnidoSite bridges the divide between high-throughput data generation and biological discovery. In summary, CnidoSite serves as a centralized and analysis-ready platform for studying core scientific questions— from coral bleaching and symbiosis breakdown to jellyfish blooms and environmental adaptation—and provides the infrastructure necessary to accelerate breakthroughs at the intersection of cnidarian biology, marine conservation, and evolutionary genomics.

## MATERIALS AND METHODS

### Genomic data collection and processing

To encompass a wide range of cnidarians, genome assembly datasets were comprehensively collected from the National Center for Biotechnology Information (NCBI) (19), Ensembl genome browser (20), Figshare (21), and the Reefgenomics database (22). The collection comprised 99 chromosome-level, 173 scaffold-level and 55 contig-level assemblies across 327 cnidarians. For species with more than one genome assembly, we retained the higher-quality, fully-annotated assembly for multi-omics analysis and visualization.

### Genome annotation and visualization

For all protein-coding genes of cnidarians, the consensus protein set was functionally annotated by sequence similarity and domain search. Proteins were searched against the NCBI non-redundant (NR) protein database (19), and the UniProt database (23) at an E-value cutoff of 1e-5, and scanned for conserved domains with InterProScan (24) using -f tsv -goterms -pa, which assigns InterPro entries, Pfam (25) and PANTHER (26) signatures together with their Gene Ontology (GO) terms (27) in a single pass. Additionally, we mapped all predicted genes to the Kyoto Encyclopedia of Genes and Genomes (KEGG) database (28) for pathway analysis. For gene family annotation, transcription factors (TFs) were classified from their conserved DNA-binding domains following AnimalTFDB 4.0 (29) with self-built HMM profiles, and ubiquitin families (Ubs) were annotated against the iUUCD database (30); both using HMMER v3.4 at a cut-off of 1e-4. Mitochondrial genomes were assembled and annotated with MitoZ (31) and MitoFinder (32) and visualized with OGDRAW v1.3.1 (33). Assembly and annotation completeness was assessed with BUSCO v5.8.3 (34) against the cnidaria_odb12 lineage (3,203 BUSCOs).

### Bulk and single-cell transcriptomic data collection and processing

Raw sequencing data were retrieved from the Sequence Read Archive (SRA), Gene Expression Omnibus (GEO), and National Genomics Data Center (NGDC) databases. Bulk transcriptome reads were quality-checked using FastQC (version 0.12.1) (35) and trimmed with Trimmomatic v0.40 (36) using default parameters, aligned to cnidarian reference genomes using HISAT2 v2.2.2 (37), and expression levels were quantified as TPM values using featureCounts v2.0.1 (38) and StringTie v2.2.1 (39). Transcript TPM values were summed by gene to form a unified expression matrix for subsequent co-expression network construction and expression heatmap visualization. For cnidarians without an annotated reference genome, transcriptomic reads were pooled and assembled de novo. Assemblies were generated with rnaSPAdes v4.3.0 (40) for 105 species and with Trinity v2.15.2 (41) for 103 species (101 primary, 2 further), giving 208 de novo assemblies; runs exceeding ∼20 Gb of usable sequence were subsampled to this cap (Trinity: --min_contig_length 200). Redundancy was reduced at the gene level rather than with CD-HIT-EST or EvidentialGene: TransDecoder v6.0.0 (42) called ORFs of ≥100 amino acids (LongOrfs -m 100) and kept the best per transcript (Predict --single_best_only), which were then collapsed to the longest ORF per Trinity gene locus (_g identifier). With the 12 species whose transcript sets came from existing genome annotations, the module covers 220 of the 327 species in CnidoSite. Each of the 220 carries a protein set annotated with Pfam, GO and KEGG as described under “Genome annotation and visualization” section. Single-cell datasets were processed as follows: 1) Data source. Count matrices and cell-type annotations were taken from the original studies; where only a processed object was available, the data were re-analysed as described below. 2) Quality control. Genes detected in fewer than three cells were discarded, and cells with fewer than 200 genes or 500 counts were removed. 3) Mitochondrial and doublet filtering. For the two datasets with a resolvable mitochondrial gene set, one was filtered at 5 MADs on mitochondrial content and the other at the 20% cut stated by the source publication; for the remaining datasets the mitochondrial gene set could not be resolved and no mitochondrial criterion was applied. A per-library 5-MAD band on library size and gene count was applied throughout, and doublets were removed with Scrublet (43). 4) Normalization and dimensionality reduction. Data were normalized, and 2,000 highly variable genes and 50 principal components were retained. 5) Batch correction, clustering and embedding. Library batch effects in multi-library datasets were corrected with Harmony (44), followed by UMAP embedding and Leiden clustering (resolution = 1.0). 6) Marker genes. Per-cluster marker genes were identified using the Wilcoxon rank-sum test.

### Proteomic data collection and processing

Publicly available proteomic datasets for cnidarians were retrieved from the PRoteomics IDEntifications (PRIDE) database (45) and the ProteomeXchange consortium (46). Acquired spectra were converted to the standard MGF format using ThermoRawFileParser (47) and searched with Comet (48) against the corresponding cnidarian protein database together with the cRAP contaminant set. Enzyme specificity and precursor and fragment mass tolerances were set per dataset to match the instrument that produced the data and, where reported, the source publication, and are recorded in each dataset’s parameter file; for the tryptic datasets this was trypsin with at most two missed cleavages, while datasets deposited as endogenous peptides were searched without enzyme specificity. Carbamidomethylation of cysteine (+57.021464) was applied as a fixed modification and oxidation of methionine (+15.9949) as a variable modification throughout, with further variable modifications added only for the datasets whose source study reported them. Peptide-spectrum matches (PSMs) were validated with Percolator (Crux 4.2) (49) against a Comet-internal reversed 1:1 decoy database at a 1% PSM and 1% peptide false discovery rate, and proteins were quantified by spectral counting. Peptides were mapped to genes through their protein identifiers.

### Epigenomic data collection and processing

Publicly available epigenomic datasets were retrieved from the Sequence Read Archive (SRA), Gene Expression Omnibus (GEO), National Genomics Data Center (NGDC) databases, and published literature. For ChIP-seq, ATAC-seq and DNase-seq, reads were aligned to the corresponding reference genome with Bowtie2 v2.5.5 (50) using “--very-sensitive --dovetail --no-mixed --no-discordant”. Enriched regions were identified with MACS2 v2.2.9.1 (51), which built the peak model from the cross-correlation of each dataset; no fixed shift or extension was applied. Because the assays differ in chemistry, peaks were called separately for each. ChIP-seq was called with “-f BAMPE” for paired-end and “-f BAM” for single-end libraries, the matched input or IgG as control, and the effective genome size of the species (e.g., 1.98e8 for *Exaiptasia diaphana*, 2.61e8 for *Nematostella vectensis*), using “--broad --broad-cutoff 0.1” for the broad histone marks H3K27me3, H3K36me3, H3K4me1 and H4K20me1 and narrow peaks for all others. ATAC-seq was called with the same genome size and control scheme, and “-f BAMPE” or “-f BAM” according to the library. DNase-seq was called with “-f BAM -g 4.38e8” (*Acropora digitifera*) and without a control. For WGBS data, adapters and low-quality bases were removed with Trimmomatic (36), and clean reads were aligned to the reference genome and deduplicated using Bismark v0.25.1 (52) over Bowtie2 v2.5.5. Gene-level methylation levels were quantified as the ratio of methylated to total reads over the gene body. For miRNA-seq data, microRNA genes, their genomic coordinates and their expression levels across developmental stages were obtained from MirGeneDB (53), with miRBase (54) identifiers used for nomenclature.

### Metagenomic data collection and processing

Metagenomic samples and microbial genomes (metagenome-assembled genomes, MAGs) derived from diverse cnidarian hosts were collected from the Gene Expression Omnibus (GEO), Genome Taxonomy Database (GTDB) (55) and published literature. Sample metadata, including host taxonomy, species identity, experimental treatments, tissue types, and developmental stages, were manually curated to ensure consistency. MAG quality was assessed using CheckM v1.1.0 (56), retaining assemblies with high completeness and low contamination. The retained MAGs were functionally annotated with KofamScan (57) and InterProScan (24).

### Co-expression network construction

Gene co-expression networks were constructed by integrating Pearson correlation coefficient (PCC) with the Mutual Rank (MR) algorithm. PCCs quantified the pairwise expression similarities across the transcriptome, while the MR metric—derived by ranking PCCs for each gene—effectively mitigated noise and prioritized robust co-expression linkages. To get optimal gene pairs and evaluate the credibility of co-expression network, candidate cut-offs (PCC > 0.7, > 0.8, > 0.9; MR top-3 with MR ≤ 30, ≤ 50 and ≤ 100) were scored against genes with known GO terms and compared by area under the ROC curve. The larger the area under the curve (AUC) value of co-expression network is, the higher the credibility of the network will be. Finally, we select the appropriate PCC and MR threshold with the highest AUC value for network construction. This dual-thresholding approach retained high-confidence interactions, which were separated into positive and negative networks (58) and visualized in the browser using Cytoscape (59).

### Phylogenetic tree construction

Genome-wide orthology inference was performed using OrthoFinder v2.5.5 (60) to delineate orthogroups (OGs). The initial phylogenomic dataset comprised 148 cnidarian species and four sponge species (phylum Porifera), *Corticium candelabrum, Halichondria panicea, Oscarella lobularis,* and *Sycon ciliatum*, with the sponge species serving as outgroups in subsequent phylogenetic analyses. To construct the phylogenomic dataset, we retained only orthogroups that contained sequences from all sampled species (i.e., 100% taxon occupancy). Within these orthogroups, when a species was represented by more than one sequence, only the longest sequence from that species was kept, so that each species was represented by exactly one sequence per orthogroup. Each OG was aligned independently with MAFFT v7.526 (61) using the ‘--anysymbol’ option, which permits non-standard amino acid characters in the input sequences. Poorly aligned and ambiguous positions were removed with BMGE v1.12 (62) under default parameters to reduce phylogenetic noise. The trimmed alignments were concatenated into a supermatrix with AMAS (63), which also generated a partition file recording the boundaries of each OG. Before phylogenetic inference, *Alatina alata* and *Calvadosia cruxmelitensis* were excluded from the concatenated alignment. The final supermatrix used for phylogenetic analyses comprised 150 species (146 cnidarian species and four sponge outgroups), 70 OGs, and 24,975 amino acid sites.

Maximum-likelihood (ML) trees were inferred from the supermatrix with IQ-TREE v3.1.2 (64) under two widely used models that account for different sources of substitution heterogeneity.

1. Partitioned model. Each OG was treated as a separate partition. ModelFinder (65) selected the best-fitting site-homogeneous substitution model for each partition independently according to the Bayesian information criterion (BIC). We applied an edge-proportional partition model (66), in which all partitions share the same topology and relative branch lengths. Each partition has its own evolutionary rate, which scales its branch lengths proportionally. The analysis was run with the options (“-s supermatrix.phy -st AA -p partition.nex -m MFP -bb 1000 --alrt 1000 --bnni”).
2. Site-heterogeneous PMSF model. To account for compositional heterogeneity across sites, we combined the LG amino acid replacement matrix (67) with the C60 empirical profile mixture model (68). C60 comprises 60 predefined amino acid equilibrium frequency profiles. Because full mixture-model inference is computationally demanding, we used the posterior mean site frequency (PMSF) approximation (69). PMSF uses a guide tree to estimate a site-specific amino acid frequency profile for each alignment position under the mixture model. We followed the two-step procedure recommended in the IQ-TREE documentation (https://iqtree.github.io/doc/iqtree-doc.pdf). A guide tree was first inferred under the LG+G4 model (“-s supermatrix.phy -st AA -m LG+G4”). The final ML tree was then inferred under the PMSF approximation of the LG+C60+G4 model using this guide tree (“-s supermatrix.phy -st AA -m LG+C60+G4 -ft guide.treefile -bb 1000 --alrt 1000 –bnni”).

Furthermore, a phylogenetic tree was reconstructed with a topology constraint derived from DeBiasse et al. (2024) (70). Reference taxa absent from our dataset were pruned, and the constraint was extended to our sampled major clades, retaining the reference backbone. The constrained topology was applied via the “-g” option in IQ-TREE, with model selection performed using ModelFinder (65) (“-s supermatrix.phy -st AA -m MFP -g constraint_tree.treefile -bb 1000 --alrt 1000 –bnni”). ModelFinder selected VT+F+I+R10 (71), which was used for the final constrained analysis.

For all aforementioned analyses, branch support was assessed with 1,000 ultrafast bootstrap replicates (UFBoot2) (72). We also ran the SH-like approximate likelihood ratio test (SH-aLRT) with 1,000 replicates (73), which provides a complementary measure of branch support. Support values are reported as SH-aLRT/UFBoot. The resulting phylogenies were visualized with phylotree.js (74).

### Database implementation

CnidoSite was constructed under a robust LAMP environment following established methodologies (75, 76). The interactive interface was implemented with HTML, CSS, JavaScript, PHP, Perl, and Python languages on Ubuntu Linux powered by an Apache server to facilitate users accessing functional genomic and multi-omics information, ensuring smooth data extraction, visualization, and presentation. CnidoSite is freely accessible to all users, without login or registration.

## DATABASE CONTENT AND USAGE

### Construction and overview of CnidoSite

To provide a comprehensive, analysis-ready multi-omics platform for Cnidaria research, CnidoSite was constructed by integrating multi-omics data from 327 cnidarians, encompassing 18,094 high-throughput sequencing samples with a total of 16TB data (Table 1). CnidoSite comprises 10 core modules: taxonomy, paleobiology, genome, transcriptome, single-cell, proteome, epigenome, metagenome, phenotype, and tools (Figure 1). The Taxonomy module retrieves taxonomy data from the NCBI Taxonomy database (77), the World Register of Marine Species (WoRMS) (78), and the Global Biodiversity Information Facility (GBIF) (79), encompassing a comprehensive taxonomy coverage including 327 species across 120 families, 23 orders, and 7 classes. Paleobiology module provides a concise overview of 411 fossil records for each taxon of cnidarians collected from the Paleobiology Database (PBDB) (80). Genome module features various genomic information, including biology descriptions, assembly details, BUSCO genes, transposable elements, gene annotation, gene families, genome phylogeny, and mitochondrial annotation. Transcriptome module collects detailed information of 15,642 transcriptomic samples across diverse adult tissues and developmental stages, providing co-expression network and dynamic network views with expression level for 29 cnidarians. Single-cell module summarizes 50 single-cell RNA-seq datasets from cnidarians to enable the identification of cell types and marker genes, and to visualize gene expression patterns across distinct cell types. Proteome module includes 46 proteomic projects with protein expression profiles in 28 species across seven classes. Epigenome module incorporates 193 ChIP-seq, 210 ATAC-seq, 1 DNase-seq, 266 WGBS, and 53 miRNA-seq datasets to comprehensively profile histone modification peaks, open chromatin regions, DNase I hypersensitive (DHS) sites, DNA methylation levels, and miRNA annotations, respectively. Metagenome module covers 1,650 metagenomic datasets from 28 projects and a corresponding set of 315 metagenome-assembled genomes (MAGs) from diverse cnidarian hosts. Phenotype module integrates 180,652 phenotypic data from multiple sources, including The Coral Trait Database (81), the Octocoral Trait Database (82), Marine Species Traits from WoRMS and the Pelagic Species Trait Database (83). Moreover, CnidoSite offers functional tools for enrichment analysis, expression visualization, BLAST alignment, primer design, and interactive visualization via GSEA, expression heatmap, BLAST, Primer3Plus (84), and JBrowse (85), respectively. CnidoSite further allows the modules to be used together through cross-links at the species and gene levels. A species portal, reachable from the Taxonomy and Genome modules and summarized as an exportable Data Coverage Matrix (327 species × 16 data types), links all data held for a species. The gene detail page in turn consolidates a gene’s genomic context, functional annotation, orthogroup and gene tree, and its multi-omics evidence, with each panel indicating data availability and linking to the corresponding viewer.

**Figure 1.**
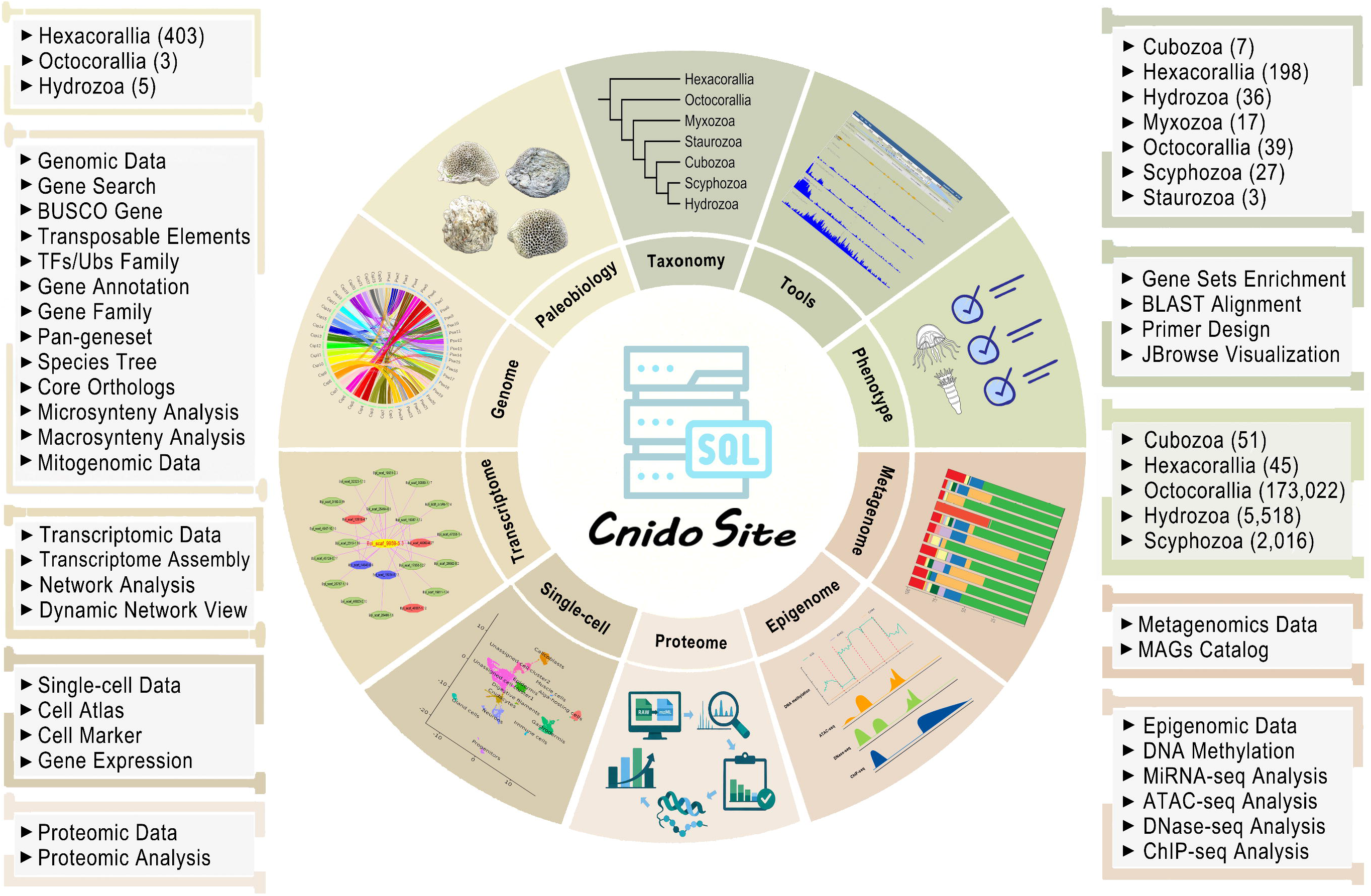
Construction and overview of CnidoSite. Database architecture and function. Distinct colors represent different modules in CnidoSite.

**Table 1.**
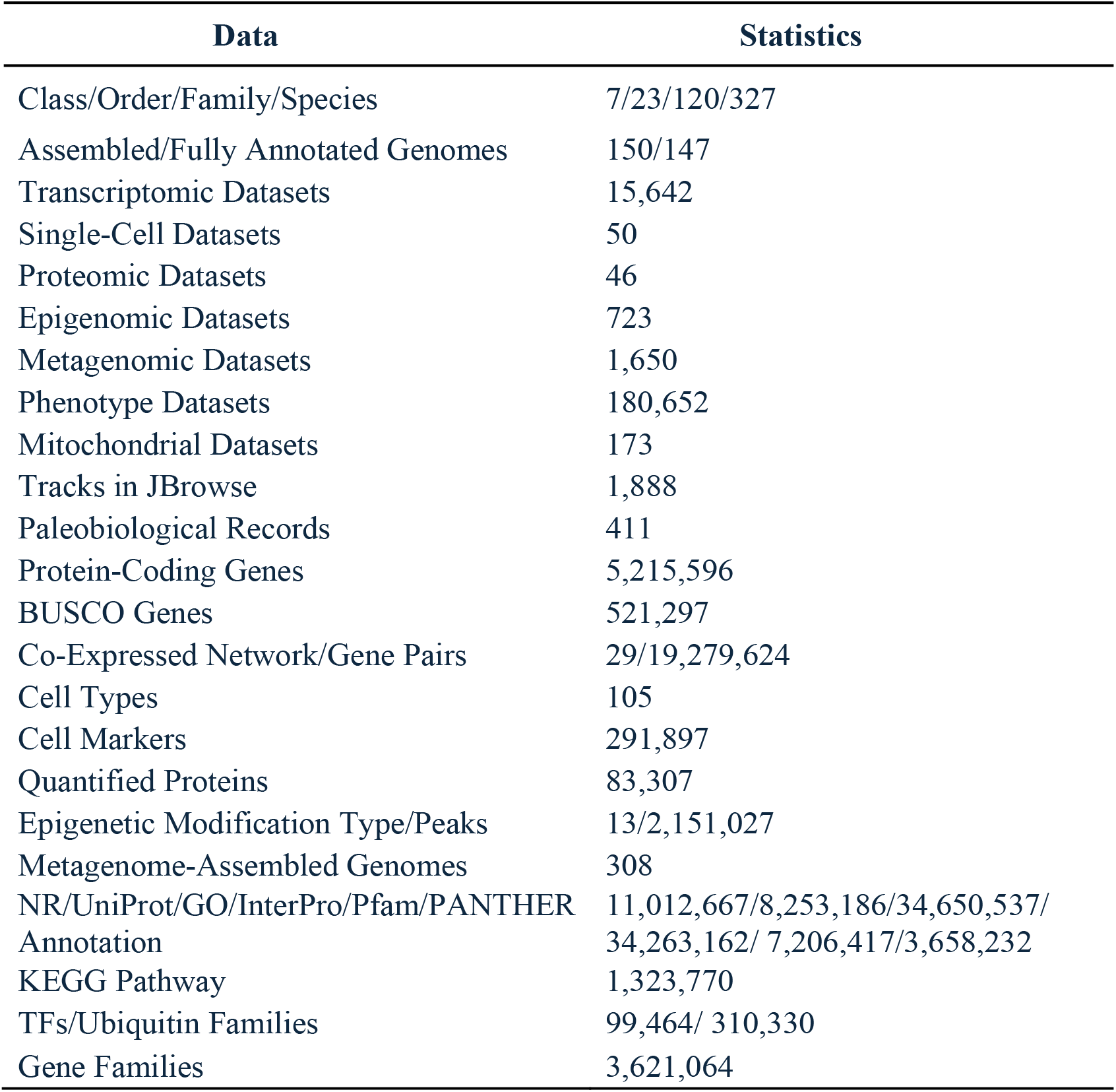
Summary of CnidoSite data composition.

### Phylogenomic backbone of Cnidaria

To trace the evolutionary lineages among these cnidarians, we inferred ML phylogenies from the supermatrix of 70 single-copy OGs (24,975 amino acid sites) under both a partitioned and a site-heterogeneous C60-PMSF model (see Methods; Figure S1A-B). Both C60-PMSF and partitioned analyses recovered monophyletic Hexacorallia, Octocorallia, Myxozoa, Cubozoa, Scyphozoa, and Hydrozoa, each with SH-aLRT/UFBoot support of 100/100. Within Medusozoa, both analyses strongly supported Cubozoa + Staurozoa (100/100), resembling the mitochondrial result of Kayal *et al*. (86), rather than the commonly recovered Cubozoa + Scyphozoa relationship. Acraspeda was recovered with lower support (70.1/76 and 60.9/74 in the C60-PMSF and partitioned analyses, respectively). Myxozoa was sister to Anthozoa + Medusozoa, contrasting with the Endocnidozoa (Myxozoa + Polypodium) – Medusozoa sister relationship commonly reported previously (70, 87–88); Polypodium was absent from our dataset. These discrepancies may partly reflect our strict requirement for 100% taxon occupancy, which excludes OGs missing from any sampled species, and alignment trimming, which removes ambiguous regions but may also discard informative sites. The resulting dataset contained 70 OGs and 24,975 amino acid sites, potentially altering the balance of signal for deep relationships compared with previous datasets. Differences in how models accommodate across-site amino acid frequency heterogeneity may also contribute, as illustrated by the model-dependent medusozoan relationships reported by DeBiasse et al. (70). Finally, Kayal *et al*. (88) reported that residual fish-host contamination distorted myxozoan placement in their preliminary analysis. If present in our input data, such non-cnidarian sequences could obscure myxozoan affinities and contribute to their placement outside Anthozoa + Medusozoa. To provide a complementary phylogenetic framework consistent with the consensus topology of major cnidarian lineages supported by previous studies, we also repeated the analysis under a constraint derived from DeBiasse *et al*. (70). The resulting constraint tree agreed with the reference backbone of DeBiasse *et al*. (70), recovering Myxozoa as sister to Medusozoa and Staurozoa as sister to Cubozoa + Scyphozoa (Figure S1C).

### Taxonomy coverage, genomic resources and paleobiology records

The phylum Cnidaria is generally categorized into seven distinct classes: Hexacorallia, Octocorallia, Myxozoa, Staurozoa, Cubozoa, Scyphozoa, and Hydrozoa. Multi-level species resources are provided in CnidoSite across all seven Cnidaria classes. At the genome level, CnidoSite incorporates 150 assembled genomes, of which 147 carry gene models (Table 2) and features a phylogenetic tree based on single-copy genes to trace the evolutionary lineages among these cnidarians on the main page. Users can select species or click on names in the “Taxonomy” module (Figure 2A) to view a brief biological introduction of each cnidarian and its genomic features (Figure 2B). Fossils provide key insights into cnidarian evolutionary history and phylogenetics (89, 90). Users can click on a taxon name in the “Paleobiology” module to reach the external PBDB database for accessing related paleobiology information, including taxonomy history, classification, morphology, and age range (Figure 2C). Owing to their high accessibility, mitochondrial genomes have emerged as a cornerstone resource for studying cnidarian evolution and phylogenetics (86, 91). We integrated 173 cnidarian mitochondrial genomes across all seven classes. Each genome is visualized as a Circos plot, and its mitochondrial gene information is provided in the “Mitogenomic Data” module (Figure 2D). Phenotype data that map trait diversity across species help to reveal the general principles that determine species distribution and abundance, evolutionary dynamics, and ecosystem functioning (82). Based on these phenotype data provided in CnidoSite, users can easily query and visualize specific trait profiles across diverse taxa in the “Phenotype” module (Figure 2E).

**Figure 2.**
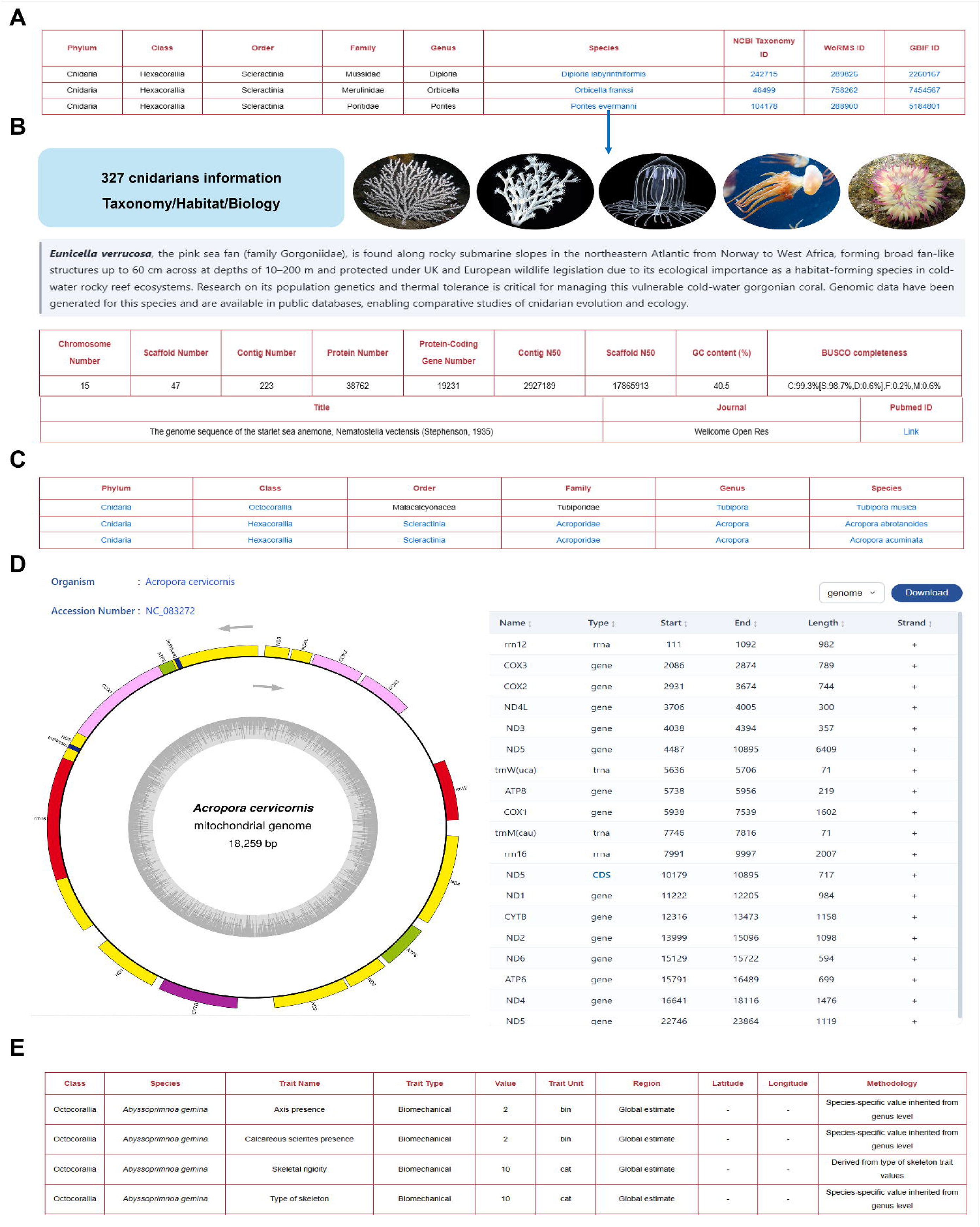
Taxonomy coverage, genomic resources and paleobiology records. Screenshots for (A) taxonomy information, (B) cnidarian information, (C) paleobiology information, (D) mitogenomic annotation, and (E) paleobiology records.

**Table 2.**
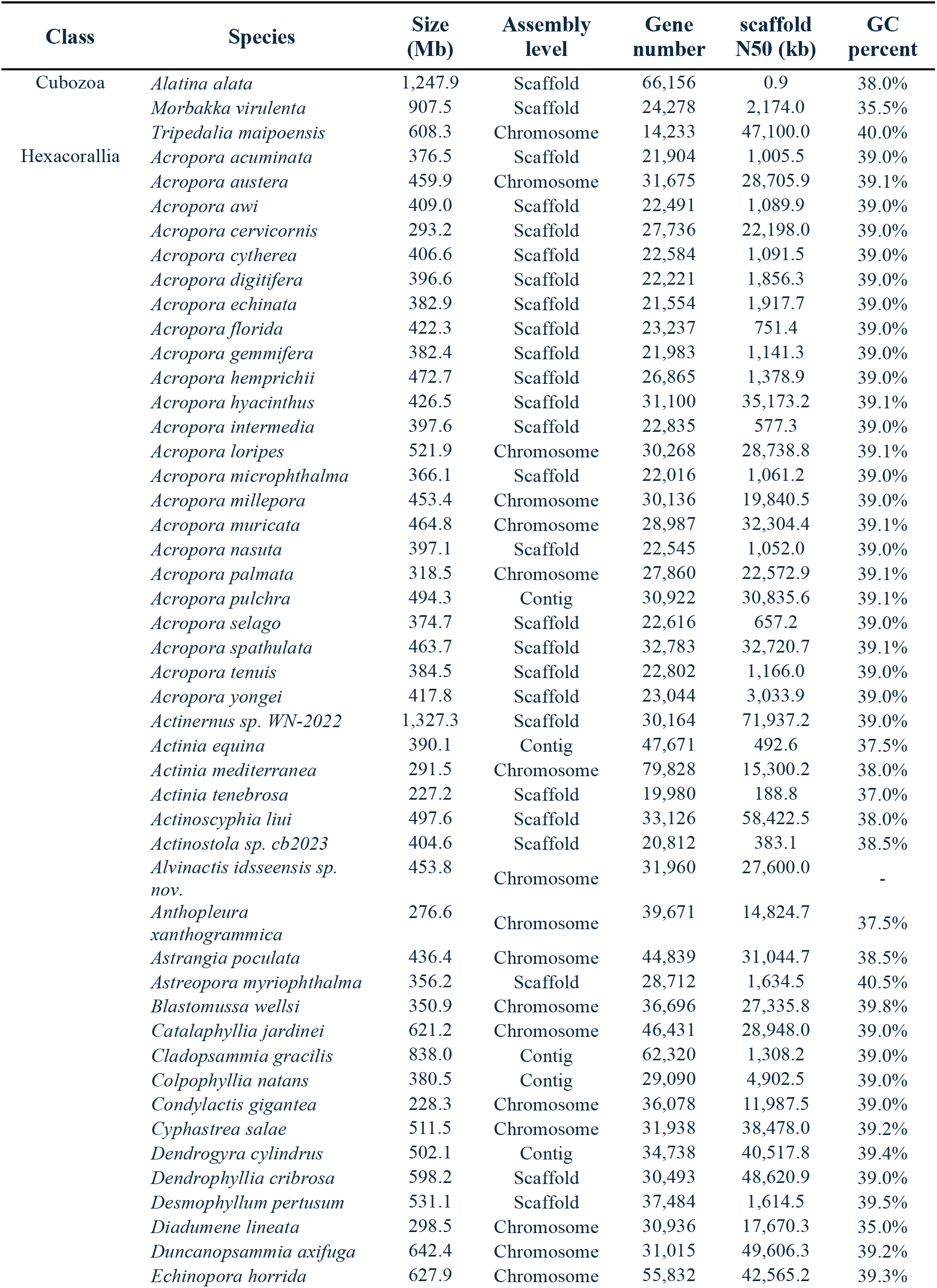
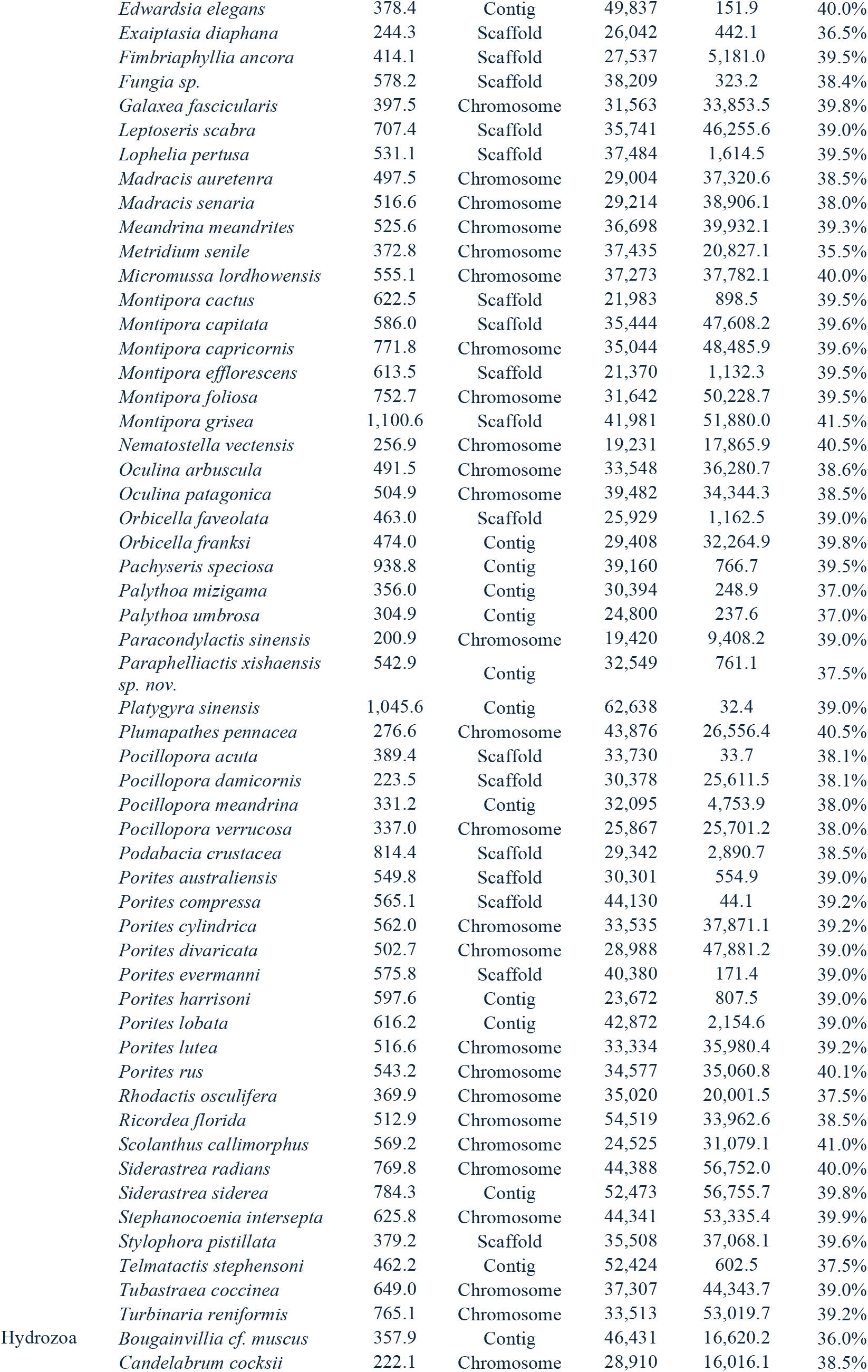
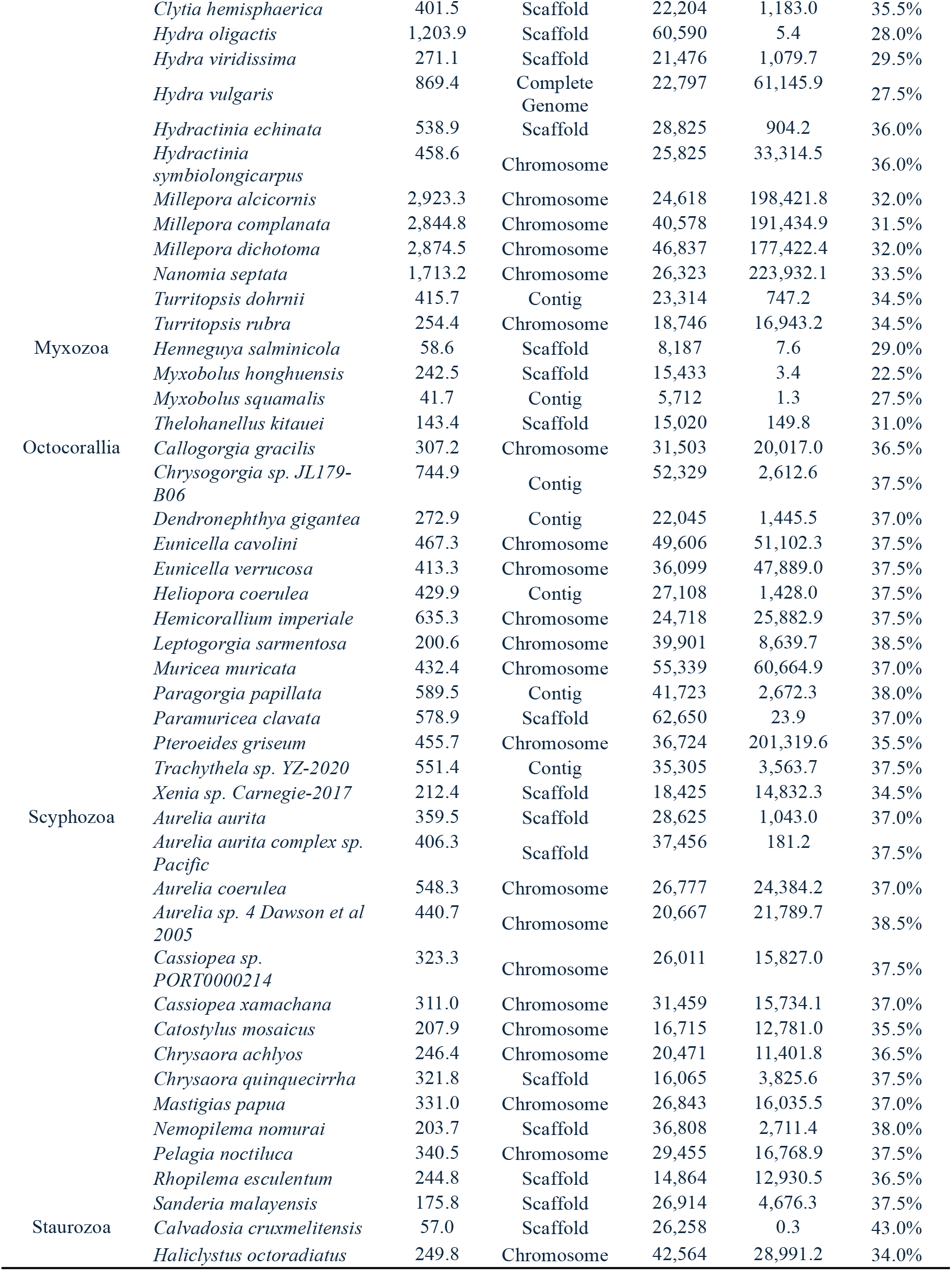
Summary of 150 cnidarian genomes with functional annotation.

### Gene functional annotation, transposable elements and gene family

For the phylum Cnidaria, functional annotation based on sequence homology is essential to bridge genomic data with physiological mechanisms, particularly in identifying genes that regulate cnidocyte development, symbiont-host interactions, and metabolic responses to climate-induced bleaching (92). A total of 5,215,596 cnidarian protein-coding genes (PCGs) were comprehensively annotated against major public databases, including NR (19), UniProt (23), Pfam (25), InterPro (24), PANTHER (26), GO (27), and KEGG (28). By searching gene IDs, genomic locus, functional keywords, or protein domains (Figure 3A) in the “Gene Search” module, users can access detailed annotation information and links to external annotation databases (Figure 3B). BUSCO gene completeness and uniqueness can provide robust, quantitative benchmarks to measure genome assembly and annotation quality, and serve as invaluable markers for phylogenetic tree construction and comparative genomics (34). A total of 521,297 BUSCO genes were identified within the predicted PCGs for each cnidarian using the cnidaria_odb12 reference database (Figure 3C). As ubiquitous features of cnidarian genomes, transposable elements (TEs) serve as pivotal evolutionary forces shaping genome plasticity, functional regulation, and adaptive evolution in these early-diverging metazoans (93). To explore the relationship between TEs and potential target genes, all annotated TEs were mapped to their nearest PCGs based on genomic distance. Users can search for a given genomic interval or TE subfamily type to view detailed annotation information (Figure 3D). Focusing on specific gene families, we classified 99,464 TF genes into 68 families, and 310,330 ubiquitin genes into 56 families. Users can search for a TF or ubiquitin family of interest in one species to obtain detailed family member information (Figure 3E-F). Basic genomic features and diverse functional elements of the 147 cnidarian genomes were visualized via customized JBrowse (Figure 3G). Users can interactively browse selected genomic regions to view comprehensive annotations, including protein sequences/structures, transposable elements (TEs), miRNA loci, and multi-omics peaks. Clicking any element opens a detailed information page, and users can upload standard-format files to create custom tracks.

**Figure 3.**
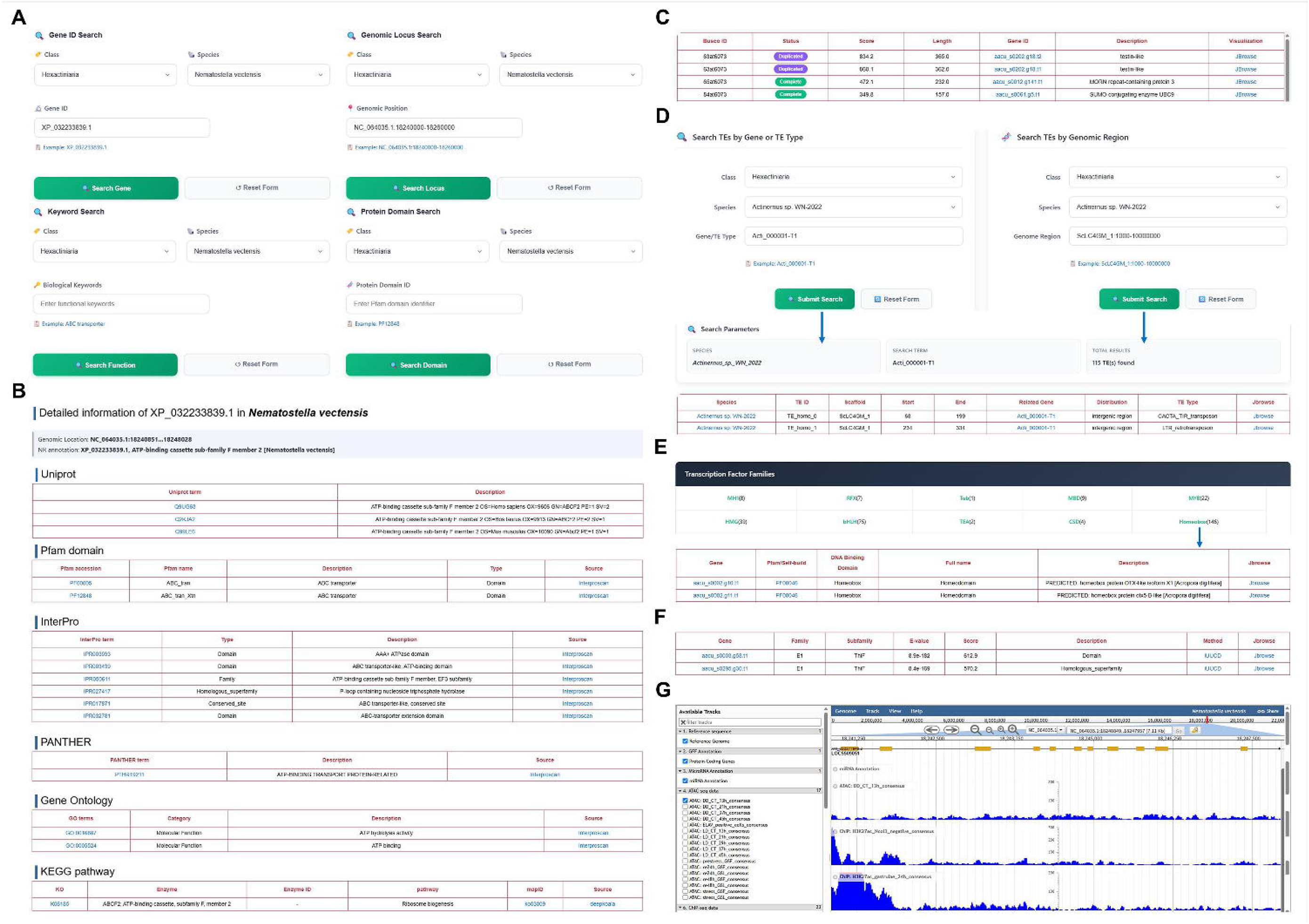
Gene functional annotation, transposable elements and gene family. Screenshots for (A) gene search, (B) gene annotation, (C) BUSCO gene, (D) transposable elements search and nearest gene information, (E) transcription factors family, (F) ubiquitin family, and (G) JBrowse visualization.

### Transcriptomic data, co-expression network visualization and analysis

Bulk transcriptomic analysis provides a comprehensive landscape of gene expression dynamics across life stages and environments, facilitating the study of evolutionary adaptation in cnidarians (94). To provide detailed information for 15,642 transcriptomic samples across all seven classes and 220 cnidarians, we manually corrected and improved major sample information, including various developmental stages or adult tissues/treatments. Users can select one species to view related transcriptomic data in the “Transcriptomic Data” module (Figure 4A). Accumulating evidence has demonstrated the efficacy of co-expression network analysis in predicting gene functions and identifying hub genes vital to distinct biological processes. Co-expression networks for 29 species were constructed based on the integration of PCC and MR algorithms. Users can select a target species and input a gene list of interest (Figure 4B) to explore the corresponding co-expression network (Figure 4C). Within the network, blue and pink lines illustrate negative and positive co-expression relationships, respectively. Additionally, Figure 4D provides detailed information regarding the co-expressed genes and their regulatory links. Based on co-expression gene pairs obtained from the preceding network, dynamic expression views with corresponding expression levels were provided to highlight the tissue- and stress-specific differentially expressed genes (Figure 4E-F). The red and blue hexagons represent up-regulated and down-regulated genes, respectively. Finally, users can perform further analysis on the network members, including expression profiling and gene set enrichment analysis (Figure 4G). The expression profiles are visualized as a heatmap, allowing users to examine gene expression patterns across various samples and identify potential tissue- and stress-specific expressions (Figure 4H). In addition, gene set enrichment analysis allows users to identify statistically significant biological processes or pathways (Figure 4I).

**Figure 4.**
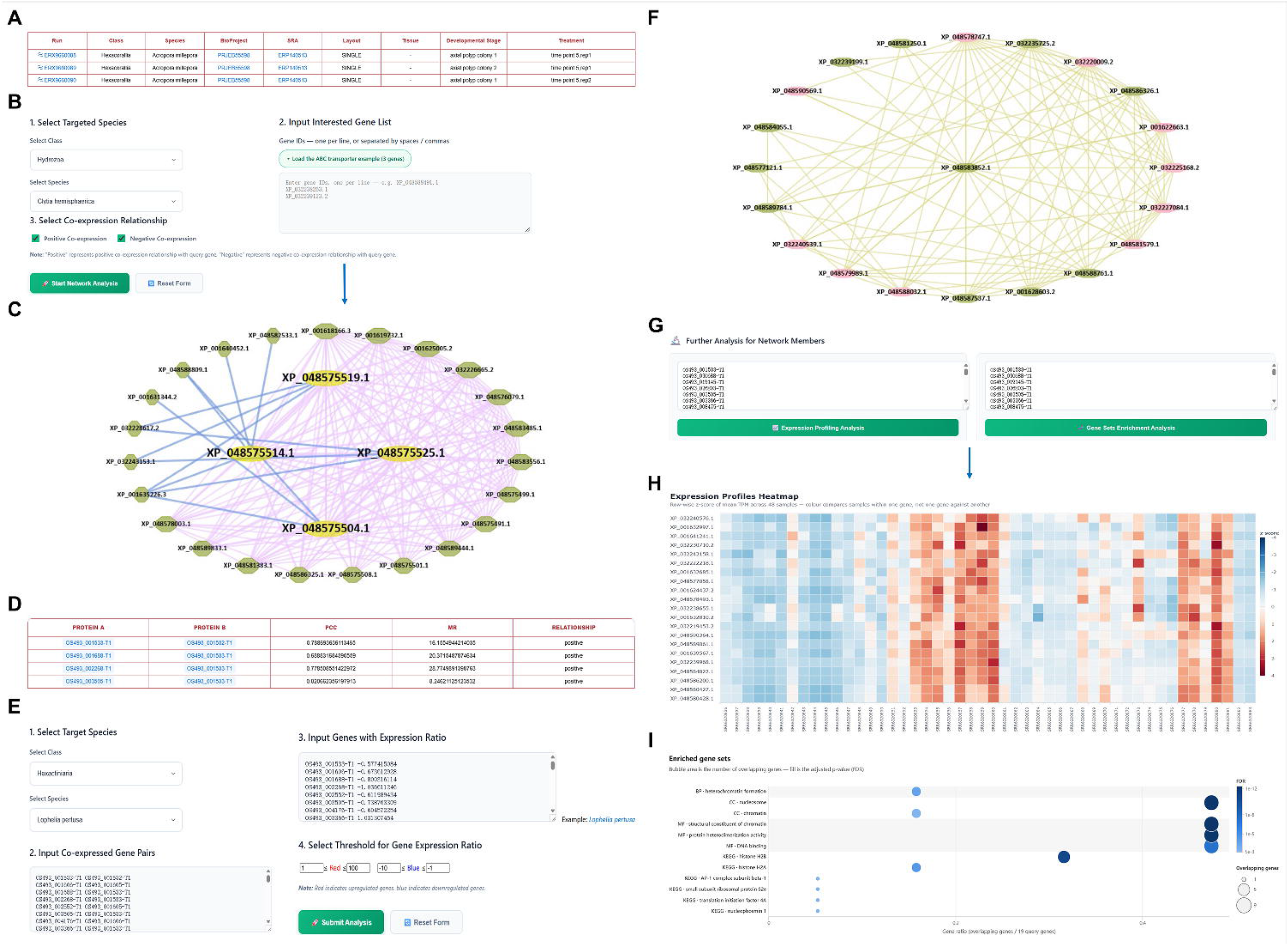
Transcriptomic data, co-expression network visualization and analysis. Screenshots for (A) transcriptomic data, (B) network search, (C) gene network visualization, (D) detailed co-expressed information, (E) dynamic network search, (F) dynamic network view, (G) further analysis for network members, (H) expression profiles, and (I) gene sets enrichment analysis results.

### Single-cell transcriptomic data visualization and marker gene analysis

Single-cell transcriptomes offer unprecedented resolution to dissect cellular heterogeneity and trace cell lineage trajectories, providing critical evolutionary insights into cell type novelties in cnidarians (15). To provide comprehensive insights into the single-cell landscape of cnidarians, we collected and uniformly processed single-cell transcriptomic data spanning major species and classes. Regarding these datasets, detailed information, including data resources, developmental stages, and cell counts, is integrated into the “Single-cell Data” module (Figure 5A). Users can select a specific tissue or developmental stage to view the corresponding cellular heterogeneity revealed via Uniform Manifold Approximation and Projection (UMAP) visualizations, presenting both unsupervised cell clusters and annotated cell types (Figure 5B). Concurrently, the “Cell Marker” module enables users to explore and analyze cell-type marker genes by specifying parameters such as species, developmental stages, and target cell types (Figure 5C-D). The top-ranked markers are displayed alongside key statistical metrics, with the option to download the complete marker gene information. Finally, a feature plot allows users to query a gene of interest and visualize its spatial expression distribution across diverse cell populations, complemented by violin plots highlighting its cell-type-specific expression dynamics (Figure 5E-F).

**Figure 5.**
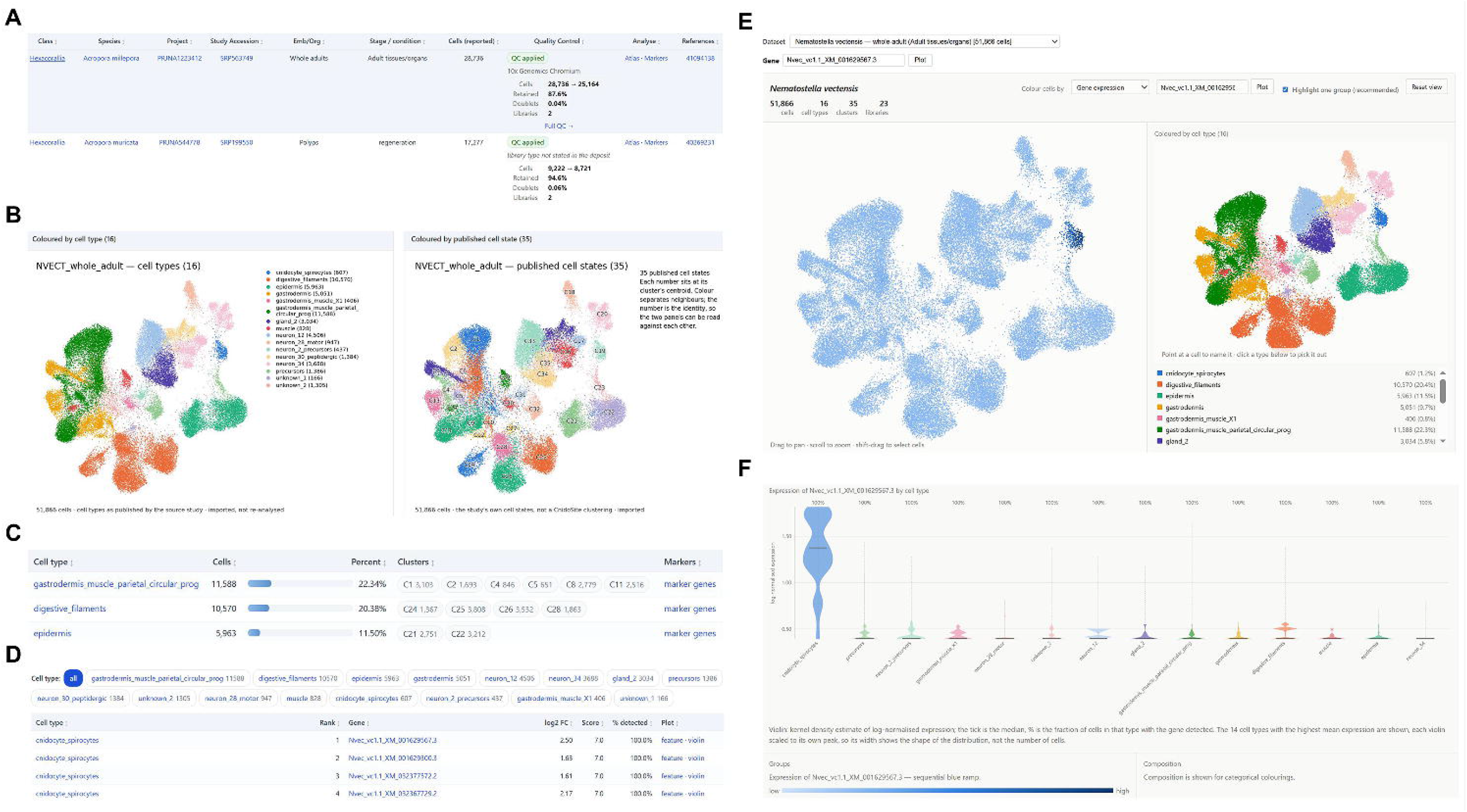
Single-cell transcriptomic data visualization and marker gene analysis. Screenshots for (A) single-cell transcriptomic data, (B) cell atlas, (C) cell marker information, (D) the expression of cell marker genes in diverse cell types, (E) gene expression across diverse cell types, and (F) violin plot for single gene expression.

### Multi-omics landscapes across proteomes, epigenomes, and metagenomes

The integration of multi-omics data provides a holistic framework linking transcriptional regulation, functional phenotypic execution, and symbiotic interactions in cnidarians (95, 96). Accordingly, a comprehensive proteomics dataset was incorporated, and users can browse metadata summaries containing classes, species, sampled tissues, experimental treatments, and corresponding primary references (Figure 6A). Upon selecting a target species, developmental stages, and tissues, users can obtain a comprehensive table displaying mass spectrometry metrics, such as identified protein IDs, unique peptide counts, sequence coverage, and high-confidence functional annotations (Figure 6B). Beyond protein profiles, users can also screen epigenomic projects based on taxonomic classes, target species, and epigenome type (Figure 6C). The interactive visualization of DNA methylation profiles allows users to analyze the genome-wide and functional-element distribution patterns of three methylation contexts (CG, CHG, and CHH). Users can query a specific gene to view nearby methylation levels and access tabular data for genome coordinates (Figure 6D). Concurrently, by querying specific miRNA-seq projects, users can retrieve detailed molecular characteristics of identified microRNAs, including miRNA ID, miRNA family, pre-miRNA or mature miRNA sequence, chromosomal coordinate, and expression level (Figure 6E). Moreover, CnidoSite integrates additional epigenomic types—including ATAC-seq, DNase-seq, and ChIP-seq (for both histone modifications and transcription factors)—which share a unified, streamlined query and visualization interface. Users can input a specific gene ID to investigate active genomic regions (Figure 6F). Related peak annotations across diverse functional regions (e.g., promoters, exons, introns, UTRs, and intergenic regions) were visualized via interactive bar and pie charts, supplemented by peak-calling statistics and genomic tracks for efficient epigenetic profiling (Figure 6F). Finally, the “Metagenome” module enables exploration of host-associated microbial communities, specifically metagenome-assembled genomes (MAGs) (Figure 6G). Users can search these MAGs by host species, accessing detailed assemblies with metrics on CheckM completeness, contamination rates, GC content, and taxonomic classifications of the symbiotic strains (Figure 6H).

**Figure 6.**
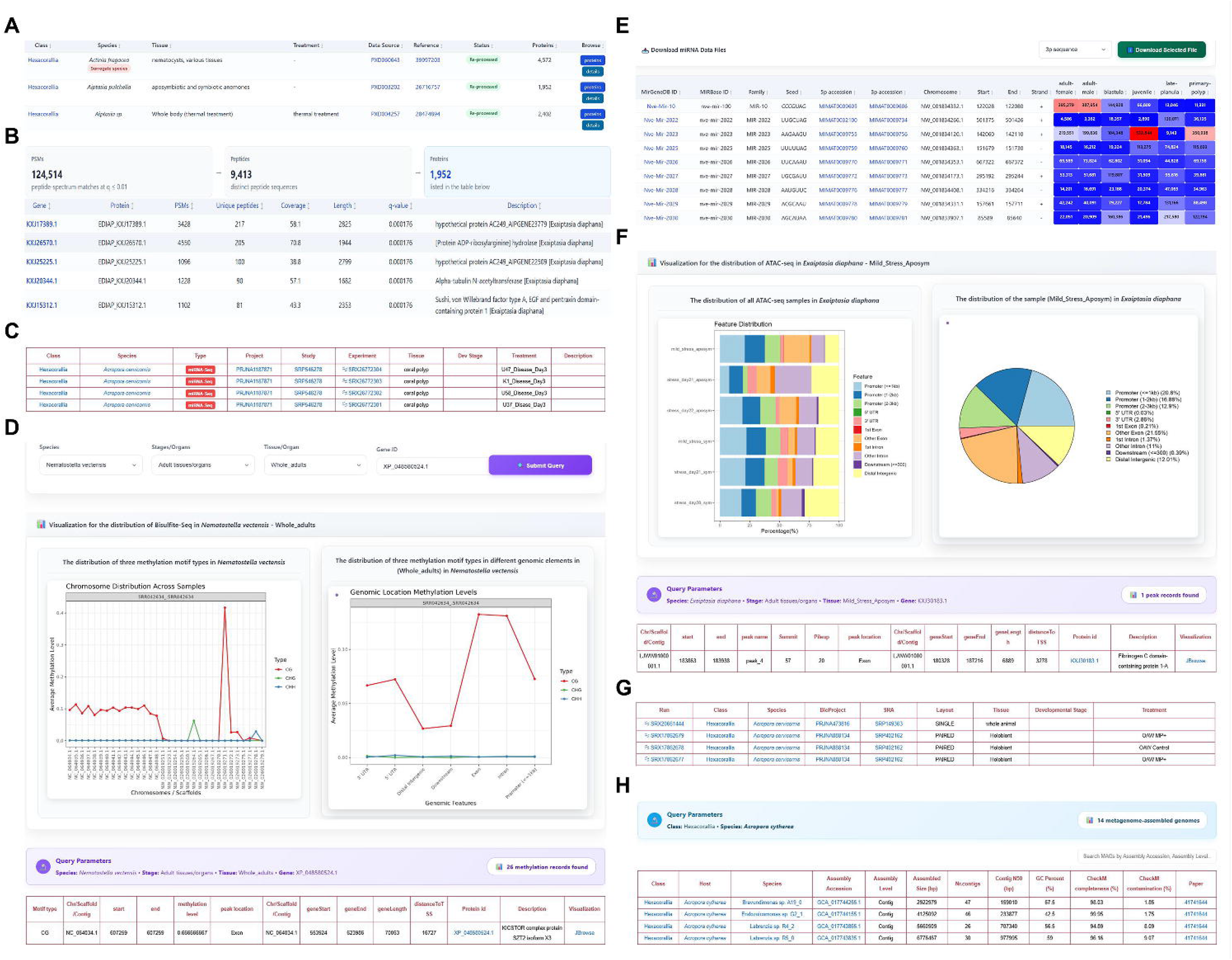
Multi-omics landscapes across proteomes, epigenomes, and metagenomes. Screenshots for (A) proteomic data, (B) proteomic analysis, (C) epigenomic search results, (D) DNA methylation analysis, (E) miRNA-seq analysis, (F) ATAC-seq, DNase-seq, and ChIP-seq analysis, (G) metagenomic data, and (H) MAGs catalog.

### Convenient online analytical tools

CnidoSite provides several interactive online tools to facilitate downstream analyses. For evolutionary studies, an integrated species tree visualization tool allows users to intuitively explore the phylogenetic relationships and taxonomic architecture of cnidarians. To decipher gene expression dynamics and functions, the platform incorporates advanced expression heatmaps alongside gene set enrichment analysis, enabling users to evaluate expression patterns across samples and pinpoint significantly enriched biological pathways. For sequence-based exploration, users can utilize BLAST alignment to search for target sequences against the reference coding sequences (CDS), proteins, or transcripts of various cnidarians. Additionally, the primer design tool allows users to specify a genomic region or directly input a sequence to generate specific primers for PCR experiments. Finally, an interactive JBrowse genomic browser is deployed, providing an intuitive, multi-track visual interface to inspect detailed genomic coordinates, gene models, and multi-omics peak annotations simultaneously.

### Future directions

CnidoSite represents a comprehensive multi-omics platform dedicated to the phylum Cnidaria, integrating genomic, bulk and single-cell transcriptomic, proteomic, epigenomic, and metagenomic resources. Although current high-quality genomic assemblies exhibit a taxonomic bias toward established model organisms, this discrepancy will gradually diminish as global sequencing initiatives expand across all cnidarian lineages. Given that cnidarian evolution, environmental adaptation, and symbiosis represent highly dynamic research frontiers, newly available, fully annotated genomes and multi-omics layers will be continuously incorporated. To circumvent current bottlenecks and maximize platform utility, we will implement regular updates centred on four strategic dimensions. First, we aim to extend taxonomic coverage by leveraging ongoing marine genome consortia to incorporate non-model species, cryptic lineages, and high-quality assembled symbiont genomes. Second, to deepen evolutionary-developmental (Evo-Devo) insights, we will integrate developmental transcriptomic datasets to enable cross-species, stage-based resolution of novel traits and life-cycle transitions. Third, every transcriptomic sample included into CnidoSite is recomputed through one pipeline against one annotation, so the pooling that makes model-based batch correction necessary does not arise. Future releases will evaluate batch-aware models wherever pooled inputs cannot be avoided. Fifth, we will extend the phylogenomic sampling to transcriptome-derived proteomes, which will allow lineages that have a genome but no gene annotation to enter the tree; the first of these will be the ceriantharian *Pachycerianthus multiplicatus*, whose position relative to Hexacorallia and Octocorallia can then be tested on data independent of genome annotation. Sixth, future releases will introduce extensive molecular marker repositories, including SNPs and microsatellites, to empower population genetics, climate resilience profiling, and genomic breeding for coral reef restoration. Finally, we will advance comparative genomics and multi-omics mining by deploying machine learning architectures and AI-driven models for automated, cross-species comparison and deep functional visualization. Together, these continuous expansions will sustain CnidoSite as an indispensable repository and interactive hub driving discoveries in cnidarian biology and marine ecosystems.

## Acknowledgements

Sincere thanks to all cnidarian researchers for access to invaluable multi-omics resources that made the work possible. We thank Ziyue Xu for kindly providing high-quality images of cnidarians featured in this study, and thank Xiyu Lin (kuroi) for her assistance in CnidoSite logo design.

## Funding

This work was funded by grants (2021HJ01, HJRC2022001, and SMSEGL26SC01) of the Southern Marine Science and Engineering Guangdong Laboratory (Guangzhou), Otto Poon Center for Climate Resilience and Sustainability (CCRS25SC01), the Research Grant Council (RGC) of the Hong Kong Special Administrative Region (26104824, 16100425, C2013-22G, and STG2/P-604/25R).

## Conflict of interest statement

None declared.

## Supplementary figure and legends

**Figure S1.**
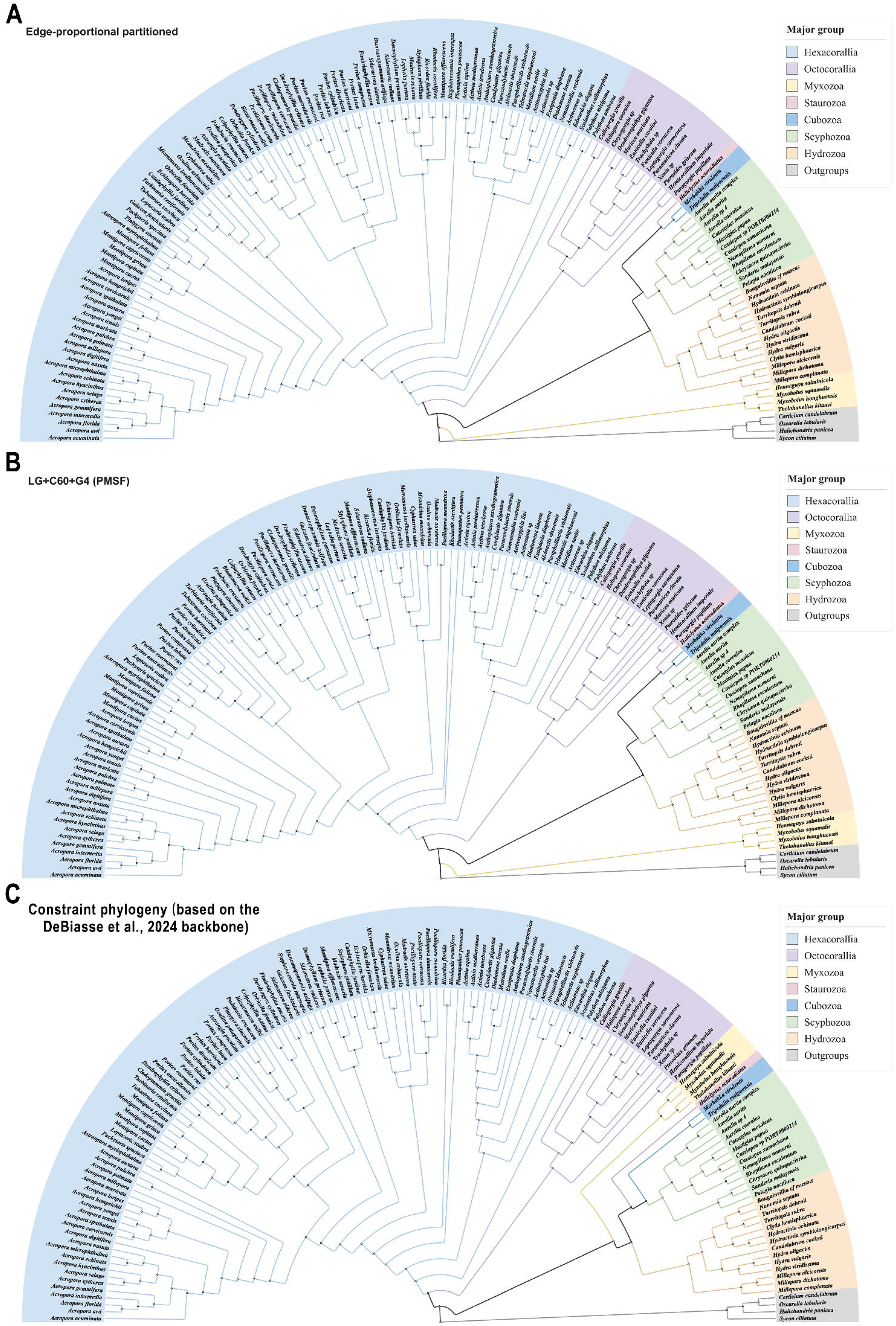
Phylogeny backbone of Cnidaria under three models. (A) Phylogeny backbone of Cnidaria under a partitioned model. (B) Phylogeny backbone of Cnidaria under LG+C60+G4 (PMSF) model. (C) Constrained phylogeny (based on the DeBiasse et al., 2024 backbone).

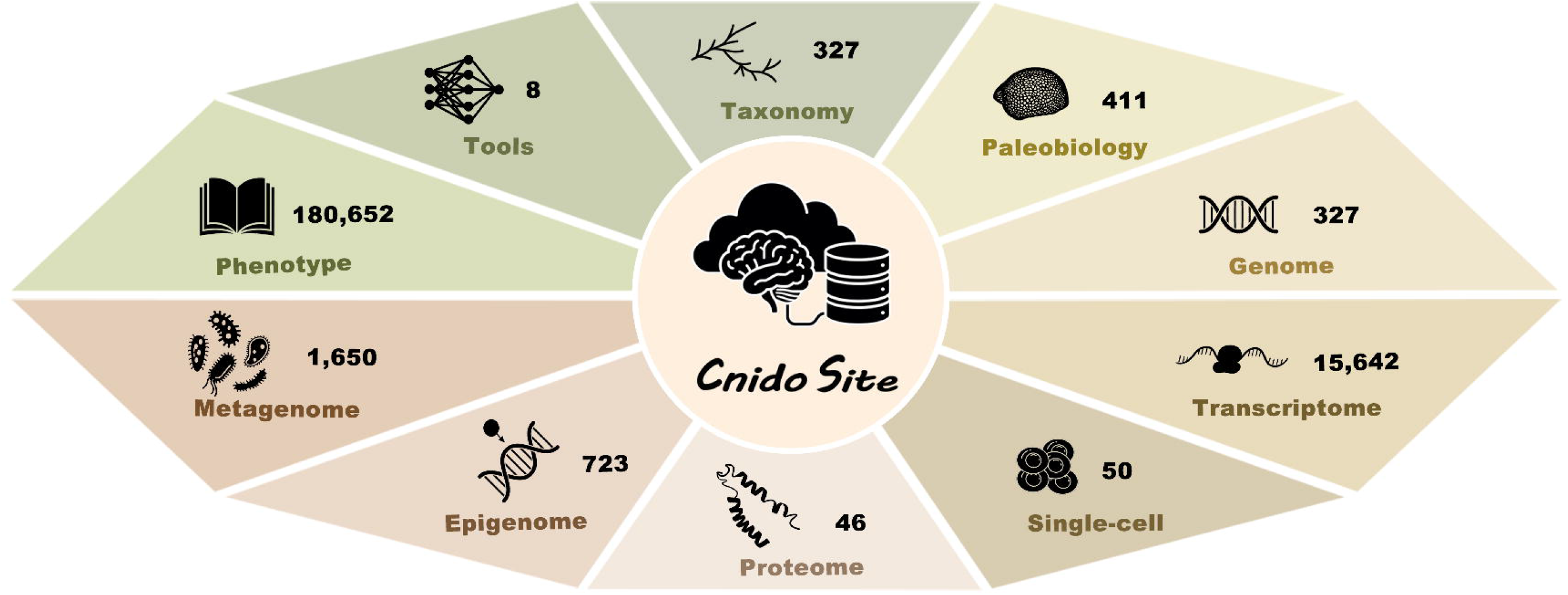

## Notes

https://cnidosite.org/

### Competing Interest Statement

The authors have declared no competing interest.

## References

1. Hoegh-Guldberg, O., Mumby, P. J., Hooten, A. J., Steneck, R. S., Greenfield, P., Gomez, E., Harvell, C. D., Sale, P. F., Edwards, A. J., Caldeira, K., et al. (2007) Coral reefs under rapid climate change and ocean acidification. Science, 318, 1737–1742.

2. Mellin, C., Brown, S., Cantin, N., Klein-Salas, E., Mouillot, D., Heron, S. F. and Fordham, D. A. (2024) Cumulative risk of future bleaching for the world’s coral reefs. Science Advances, 10, eadn9660.

3. Hughes, T. P., Kerry, J. T., Baird, A. H., Connolly, S. R., Dietzel, A., Eakin, C. M., Heron, S. F., Hoey, A. S., Hoogenboom, M. O., Liu, G., et al. (2018) Global warming transforms coral reef assemblages. Nature, 556, 492–496.

4. Hughes T. P., Kerry J. T., Álvarez-Noriega M., et al. (2017) Global warming and recurrent mass bleaching of corals. Nature, 543, 373–377.

5. Vaga C. F., Quattrini A. M., Galvão de Lossio e Seiblitz I., et al. (2025) A global coral phylogeny reveals resilience and vulnerability through deep time. Nature, 1-6.

6. Richardson, A. J., Bakun, A., Hays, G. C., & Gibbons, M. J. (2009) The jellyfish joyride: causes, consequences and management responses to a more gelatinous future. Trends in ecology & evolution, 24, 312–322.

7. Plessier, F. and Marlow, H. (2026) Cellular and transcriptional trajectories of neural fate specification in sea anemone uncover two modes of adult neurogenesis. Nature Communications, 17, 2611.

8. Sebé-Pedrós A., Chomsky E., Pang K., et al. (2018) Early metazoan cell type diversity and the evolution of multicellular gene regulation. Nature ecology & evolution, 2, 1176–1188.

9. Elek, A., Iglesias, M., Mahieu, L., Zolotarov, G., Grau-Bové, X., Aerts, S. and Sebé-Pedrós, A. (2026) Decoding cnidarian cell type gene regulation. Nature Ecology & Evolution, 10, 140–153.

10. Kim, I.V., Navarrete, C., Grau-Bové, X., Iglesias, M., Elek, A., Zolotarov, G., Bykov, N.S., Montgomery, S.A., Ksiezopolska, E., Cañas-Armenteros, D., et al. (2025) Chromatin loops are an ancestral hallmark of the animal regulatory genome. Nature, 642, 1097–1105.

11. Putnam N. H., Srivastava M., Hellsten U., et al. (2007) Sea anemone genome reveals ancestral eumetazoan gene repertoire and genomic organization. Science, 317, 86– 94.

12. Hughes, T.P., Barnes, M.L., Bellwood, D.R., Cinner, J.E., Cumming, G.S., Jackson, J.B.C., Kleypas, J., van de Leemput, I.A., Lough, J.M., Morrison, T.H., et al. (2017) Coral reefs in the Anthropocene. Nature, 546, 82–90.

13. Nong, W., Cao, J., Li, Y., Qu, Z., Sun, J., Swale, T., Yip, H.Y., Qian, P.Y., Qiu, J.-W., Kwan, H.S., et al. (2020) Jellyfish genomes reveal distinct homeobox gene clusters and conservation of small RNA processing. Nature Communications, 11, 3051.

14. Cole A. G., Steger J., Hagauer J., et al. (2024) Updated single cell reference atlas for the starlet anemone *Nematostella vectensis*. Frontiers in Zoology, 21, 8.

15. Levy S., Elek A., Grau-Bové X., et al. (2021) A stony coral cell atlas illuminates the molecular and cellular basis of coral symbiosis, calcification, and immunity. Cell, 184, 2973–2987.

16. Ryan, J. F., Finnerty, J. R. (2003) CnidBase: The cnidarian evolutionary genomics database. Nucleic Acids Research, 31, 159–163.

17. Voolstra C. R., Quigley K. M., Davies S. W., et al. (2021) Consensus guidelines for advancing coral holobiont genome and specimen voucher deposition. Frontiers in Marine Science, 8, 701784.

18. Bao Y., Song S., He S., et al. (2025) Database resources of the National Genomics Data Center, China National Center for Bioinformation in 2026. Nucleic Acids Res, 54, D28–D47.

19. Sayers, E.W., Beck, J., Bolton, E.E., Brister, J.R., Chan, J., Connor, R., Feldgarden, M., Fine, A.M., Funk, K., Hoffman, J., et al. (2024) Database resources of the National Center for Biotechnology Information in 2025. Nucleic Acids Res, 53, D20– D29.

20. Yates A. D., Austine-Orimoloye O., Azov A. G., et al. (2026) Ensembl 2026. Nucleic Acids Res, 54, D1053–D1060.

21. Singh, J. (2011) FigShare. J Pharmacol Pharmacother, 2, 138–139.

22. Liew Y. J., Aranda M., Voolstra C. R. (2016) Reefgenomics. Org-a repository for marine genomics data. Database, 2016, baw152.

23. The UniProt Consortium (2021) UniProt: the universal protein knowledgebase in 2021. Nucleic Acids Research, 49, D480–D489.

24. Blum, M., Andreeva, A., Florentino, L.C., Chuguransky, S.R., Grego, T., Hobbs, E., Pinto, B.L., Orr, A., Paysan-Lafosse, T., Ponamareva, I., et al. (2025) InterPro: the protein sequence classification resource in 2025. Nucleic Acids Res, 53, D444–D456.

25. Mistry, J., Chuguransky, S., Williams, L., Qureshi, M., Salazar, G.A., Sonnhammer, E.L.L., Tosatto, S.C.E., Paladin, L., Raj, S., Richardson, L.J., et al. (2021) Pfam: The protein families database in 2021. Nucleic Acids Res, 49, D412–D419.

26. Mi, H., Huang, X., Muruganujan, A., Tang, H., Mills, C., Kang, D. and Thomas, P.D. (2017) PANTHER version 11: expanded annotation data from Gene Ontology and Reactome pathways, and data analysis tool enhancements. Nucleic Acids Res, 45, D183–D189.

27. Aleksander S. A., Balhoff J. P., Carbon S., et al. (2025) The Gene Ontology knowledgebase in 2026. Nucleic Acids Res, 54, gkaf1292.

28. Kanehisa, M., Furumichi, M., Sato, Y., Matsuura, Y. and Ishiguro-Watanabe, M. (2025) KEGG: biological systems database as a model of the real world. Nucleic Acids Res, 53, D672–D677.

29. Shen, W. K., Chen, S. Y., Gan, Z. Q., Zhang, Y. Z., Yue, T., Chen, M. M., Xue, Y., Hu, H. and Guo, A. Y. (2023) AnimalTFDB 4.0: a comprehensive animal transcription factor database updated with variation and expression annotations. Nucleic Acids Res, 51, D39–D45.

30. Zhou, J., Xu, Y., Lin, S., Guo, Y., Deng, W., Zhang, Y., Guo, A. and Xue, Y. (2018) iUUCD 2.0: an update with rich annotations for ubiquitin and ubiquitin-like conjugations. Nucleic Acids Res, 46, D447–D453.

31. Meng G, Li Y, Yang C, et al. (2019) MitoZ: a toolkit for animal mitochondrial genome assembly, annotation and visualization. Nucleic acids research, 47, e63–e63.

32. Allio R, Schomaker-Bastos A, Romiguier J, et al. (2020) MitoFinder: Efficient automated large - scale extraction of mitogenomic data in target enrichment phylogenomics. Molecular ecology resources, 20, 892–905.

33. Greiner, S., Lehwark, P. and Bock, R. (2019) OrganellarGenomeDRAW (OGDRAW) version 1.3.1: expanded toolkit for the graphical visualization of organellar genomes. Nucleic Acids Res, 47, W59–W64.

34. Tegenfeldt, F., Kuznetsov, D., Manni, M., Berkeley, M., Zdobnov, E.M. and Kriventseva, E.V. (2025) OrthoDB and BUSCO update: annotation of orthologs with wider sampling of genomes. Nucleic Acids Res, 53, D516–D522.

35. Andrews, S. (2010) FastQC: a quality control tool for high throughput sequence data. Available from: https://www.bioinformatics.babraham.ac.uk/projects/fastqc/ (accessed 2 October 2026).

36. Beier, S., Bolger, A. M., Bolger, M. E., Schwacke, R. and Usadel, B. (2026) Trimmomatic: a decade of feature-rich, high-performance NGS read preprocessing. Bioinformatics, 42, btag331.

37. Kim, D., Paggi, J. M., Park, C., Bennett, C. and Salzberg, S. L. (2019) Graph-based genome alignment and genotyping with HISAT2 and HISAT-genotype. Nature Biotechnology, 37, 907–915.

38. Liao, Y., Smyth, G. K. and Shi, W. (2014) featureCounts: an efficient general purpose program for assigning sequence reads to genomic features. Bioinformatics, 30, 923– 930.

39. Pertea, M., Pertea, G. M., Antonescu, C. M., Chang, T. C., Mendell, J. T. and Salzberg, S. L. (2015) StringTie enables improved reconstruction of a transcriptome from RNA-seq reads. Nature biotechnology, 33, 290–295.

40. Bushmanova E, Antipov D, Lapidus A, et al. (2019) rnaSPAdes: a de novo transcriptome assembler and its application to RNA-Seq data. GigaScience, 8, giz100.

41. Grabherr MG, Haas BJ, Yassour M, et al. (2011) Full-length transcriptome assembly from RNA-Seq data without a reference genome. Nature Biotechnology, 29, 644– 652.

42. Haas BJ, Papanicolaou A. (2016) TransDecoder v6.0.0. https://github.com/TransDecoder/TransDecoder.

43. Wolock, S. L., Lopez, R., & Klein, A. M. (2019). Scrublet: computational identification of cell doublets in single-cell transcriptomic data. Cell systems, 8, 281–291.

44. Korsunsky I, Millard N, Fan J, et al. (2019) Fast, sensitive and accurate integration of single-cell data with Harmony. Nature methods, 16, 1289–1296.

45. Perez-Riverol, Y., Bandla, C., Kundu, D. J., Kamatchinathan, S., Bai, J., Hewapathirana, S., John, N. S., Prakash, A., Walzer, M., Wang, S., et al. (2024) The PRIDE database at 20 years: 2025 update. Nucleic Acids Res, 53, D543–D553.

46. Deutsch, E. W., Bandeira, N., Perez-Riverol, Y., Sharma, V., Carver, J. J., Mendoza, L., Kundu, D. J., Bandla, C., Kamatchinathan, S., Hewapathirana, S., et al. (2026) The ProteomeXchange consortium in 2026: making proteomics data FAIR. Nucleic Acids Res, 54, D459–D469.

47. Hulstaert N, Shofstahl J, Sachsenberg T, et al. (2020) ThermoRawFileParser: modular, scalable, and cross-platform RAW file conversion. Journal of proteome research, 19, 537–542.

48. Eng, J. K., Jahan, T. A., & Hoopmann, M. R. (2013) Comet: an open-source MS/MS sequence database search tool. Proteomics, 13, 22–24.

49. The M, MacCoss M. J., Noble W. S., et al. (2016) Fast and accurate protein false discovery rates on large-scale proteomics data sets with percolator 3.0. Journal of the American Society for Mass Spectrometry, 27, 1719–1727.

50. Langmead, B. and Salzberg, S. L. (2012) Fast gapped-read alignment with Bowtie 2. Nature methods, 9, 357–359.

51. Feng J, Liu T, Qin B, et al. (2012) Identifying ChIP-seq enrichment using MACS. Nature protocols, 7,1728–1740.

52. Krueger, F. and Andrews, S. R. (2011) Bismark: a flexible aligner and methylation caller for Bisulfite-Seq applications. Bioinformatics, 27, 1571–1572.

53. Clarke, A. W., Høye, E., Hembrom, A. A., Paynter, V. M., Vinther, J., Wyrożemski, Ł., Biryukova, I., Formaggioni, A., Ovchinnikov, V., Herlyn, H., et al. (2025) MirGeneDB 3.0: improved taxonomic sampling, uniform nomenclature of novel conserved microRNA families and updated covariance models. Nucleic Acids Res, 53, D116–D128.

54. Kozomara, A., Birgaoanu, M. and Griffiths-Jones, S. (2019) miRBase: from microRNA sequences to function. Nucleic Acids Research, 47, D155–D162.

55. Parks, D. H., Chaumeil, P. A., Mussig, A. J., Rinke, C., Chuvochina, M. and Hugenholtz, P. (2026) GTDB release 10: a complete and systematic taxonomy for 715 230 bacterial and 17 245 archaeal genomes. Nucleic Acids Res, 54, D743–D754.

56. Chklovski, A., Parks, D. H., Woodcroft, B. J. and Tyson, G. W. (2023) CheckM2: a rapid, scalable and accurate tool for assessing microbial genome quality using machine learning. Nature Methods, 20, 1203–1212.

57. Aramaki T, Blanc-Mathieu R, Endo H, et al. (2020) KofamKOALA: KEGG Ortholog assignment based on profile HMM and adaptive score threshold. Bioinformatics, 36, 2251–2252.

58. Lim, P. K., Wang, R., Lim, S. C., Antony Velankanni, J. P. and Mutwil, M. (2026) Constructing gene co-functional and co-regulatory networks from public transcriptomes using condition-specific ensemble co-expression. Nature Communications, 17, 5906.

59. Otasek, D., Morris, J. H., Bouças, J., Pico, A. R. and Demchak, B. (2019) Cytoscape Automation: empowering workflow-based network analysis. Genome Biology, 20, 185.

60. Emms, D. M., Liu, Y., Belcher, L., Holmes, J. and Kelly, S. (2026) OrthoFinder: improved phylogenetic orthology inference with enhanced accuracy and scalability. Nature Methods, 23, 1327–1333.

61. Katoh, K., Kuma, K., Toh, H. and Miyata, T. (2005) MAFFT version 5: improvement in accuracy of multiple sequence alignment. Nucleic Acids Res, 33, 511–518.

62. Criscuolo, A. and Gribaldo, S. (2010) BMGE (Block Mapping and Gathering with Entropy): a new software for selection of phylogenetic informative regions from multiple sequence alignments. BMC Evolutionary Biology, 10, 210.

63. Borowiec, M. L. (2016) AMAS: a fast tool for alignment manipulation and computing of summary statistics. PeerJ, 4, e1660.

64. Wong, T. K. F., Ly-Trong, N., Ren, H., Demotte, P., Baños, H., Roger, A. J., Susko, E., Bielow, C., De Maio, N., Goldman, N., et al. (2026) IQ-TREE 3: phylogenomic inference software using complex evolutionary models. Molecular Biology and Evolution, 43, msag117.

65. Kalyaanamoorthy S, Minh B. Q., Wong T. K. F., et al. (2017) ModelFinder: fast model selection for accurate phylogenetic estimates. Nature methods, 14, 587–589.

66. Chernomor O, Von Haeseler A, Minh B. Q. (2016) Terrace aware data structure for phylogenomic inference from supermatrice. Systematic biology, 65, 997–1008.

67. Le S. Q., Gascuel O. (2008) An improved general amino acid replacement matrix. Molecular Biology and Evolution, 25, 1307–1320.

68. Si Quang L, Gascuel O, Lartillot N. (2008) Empirical profile mixture models for phylogenetic reconstruction. Bioinformatics, 24, 2317–2323.

69. Wang H C, Minh B Q, Susko E, et al. (2018) Modeling site heterogeneity with posterior mean site frequency profiles accelerates accurate phylogenomic estimation. Systematic biology, 67, 216–235.

70. DeBiasse, M. B., Buckenmeyer, A., Macrander, J., et al. (2024) A cnidarian phylogenomic tree fitted with hundreds of 18S leaves. Bulletin of the Society of Systematic Biologists, 3.

71. Müller T, Vingron M. (2000) Modeling amino acid replacement. Journal of Computational Biology, 7, 761–776.

72. Hoang D T, Chernomor O, Von Haeseler A, et al. (2018) UFBoot2: improving the ultrafast bootstrap approximation. Molecular Biology and Evolution, 35, 518–522.

73. Guindon S, Dufayard J F, Lefort V, et al. (2010) New algorithms and methods to estimate maximum-likelihood phylogenies: assessing the performance of PhyML 3.0. Systematic biology, 59, 307–321.

74. Shank, S. D., Weaver, S. and Kosakovsky Pond, S. L. (2018) phylotree.js - a JavaScript library for application development and interactive data visualization in phylogenetics. BMC Bioinformatics, 19, 276.

75. Wang, G., Schmidt, C., Li, X. and Wang, Z. (2026) APD6: the antimicrobial peptide database is expanded to promote research and development by deploying an unprecedented information pipeline. Nucleic Acids Res, 54, D363–D374.

76. Kemmler, E., Lemfack, M. C., Goede, A., Gallo, K., Toguem, S. M. T., Ahmed, W., Millberg, I., Preissner, S., Piechulla, B. and Preissner, R. (2025) mVOC 4.0: a database of microbial volatiles. Nucleic Acids Res, 53, D1692–D1696.

77. Cox, E., Tsuchiya, M. T. N., Ciufo, S., Torcivia, J., Falk, R., Anderson, W. R., Holmes, J. B., Hem, V., Breen, L., Davis, E., et al. (2025) NCBI taxonomy: enhanced access via NCBI datasets. Nucleic Acids Res, 53, D1711–D1715.

78. Ahyong, S. T., Boyko, C. B., Bernot, J., et al. (2025) World Register of Marine Species (WoRMS). WoRMS Editorial Board. Available from: https://www.marinespecies.org (accessed 2 October 2026).

79. GBIF.org. (2026) GBIF Home Page. Global Biodiversity Information Facility. Available from: https://www.gbif.org (accessed 2 October 2026).

80. Peters, S. E., & McClennen, M. (2016). The Paleobiology database application programming interface. Paleobiology, 42, 1–7.

81. Madin, J. S., Anderson, K. D., Andreasen, M. H., Bridge, T. C. L., Cairns, S. D., Connolly, S. R., Darling, E. S., Diaz, M., Falster, D. S., Franklin, E. C., et al. (2016) The Coral Trait Database, a curated database of trait information for coral species from the global oceans. Scientific Data, 3, 160017.

82. Gómez-Gras, D., Linares, C., Viladrich, N., Zentner, Y., Grinyó, J., Gori, A., McFadden, C. S., Fabricius, K. E. and Madin, J. S. (2025) The Octocoral Trait Database: a global database of trait information for octocoral species. Scientific Data, 12, 82.

83. Gleiber, M. R., Hardy, N. A., Roote, Z., Krug-MacLeod, A. M., Morganson, C. J., Tandy, Z., George, I., Matuch, C., Brookson, C. B., Daly, E. A., et al. (2024) The Pelagic Species Trait Database, an open data resource to support trait-based ocean research. Scientific Data, 11, 2.

84. Hung, J. H., & Weng, Z. (2016). Designing polymerase chain reaction primers using Primer3Plus. Cold Spring Harbor Protocols, 2016, pdb-prot093096.

85. Diesh, C., Stevens, G. J., Xie, P., De Jesus Martinez, T., Hershberg, E. A., Leung, A., Guo, E., Dider, S., Zhang, J., Bridge, C., et al. (2023) JBrowse 2: a modular genome browser with views of synteny and structural variation. Genome Biology, 24, 74.

86. Kayal E, Roure B, Philippe H, et al. (2013) Cnidarian phylogenetic relationships as revealed by mitogenomics. BMC evolutionary biology, 13, 5.

87. Chang E S, Neuhof M, Rubinstein N D, et al. (2015) Genomic insights into the evolutionary origin of Myxozoa within Cnidaria. Proceedings of the National Academy of Sciences, 112, 14912–14917.

88. Kayal E, Bentlage B, Sabrina Pankey M, et al. (2018) Phylogenomics provides a robust topology of the major cnidarian lineages and insights on the origins of key organismal traits. BMC evolutionary biology, 18, 68.

89. Mulvey, L. P., Nikolic, M. C., Allen, B. J., Heath, T. A., & Warnock, R. C. (2025). From fossils to phylogenies: exploring the integration of paleontological data into Bayesian phylogenetic inference. Paleobiology, 51, 214–236.

90. do Rosario Petrucci, B., May, M. R., & Heath, T. A. (2025). Fossils improve extinction-rate estimates under state-dependent diversification models. Philosophical Transactions of the Royal Society B: Biological Sciences, 380, 20230313.

91. Feng, H., Lv, S., Li, R., Shi, J., Wang, J. and Cao, P. (2023) Mitochondrial genome comparison reveals the evolution of cnidarians. Ecology and Evolution, 13, e10157.

92. Metz, S., Paulini, M., Rising, K., Quigley, K. M., Voolstra, C. R., Combosch, D. J., Ziegler, M., Parkinson, J. E., McKenna, V., O’Brien, R., et al. (2026) Chromosome-level genomes of scleractinian corals: gene prediction and functional annotation. Scientific Data.

93. Wong, W. Y., Simakov, O., Bridge, D. M., Cartwright, P., Bellantuono, A. J., Kuhn, A., Holstein, T. W., David, C. N., Steele, R. E. and Martínez, D. E. (2019) Expansion of a single transposable element family is associated with genome-size increase and radiation in the genus Hydra. Proc Natl Acad Sci U S A, 116, 22915–22917.

94. Wang J., Zhang L., Lian S., et al. (2020) Evolutionary transcriptomics of metazoan biphasic life cycle supports a single intercalation origin of metazoan larvae. Nature Ecology & Evolution, 4, 725–736.

95. Bhattacharya D., Nandi S., Chille EE., et al. (2026) The host coral bleaching response viewed through the lens of multi-omics: multi-omics provides the tools to understand the complex molecular basis of coral bleaching, which can aid conservation efforts. BioEssays, 48, e70110.

96. Sun, H., Liu, X., Ahmad, M., Guo, X., Yu, Z., Zhao, Y., Shen, P., Zhang, Y. and Dong, Z. (2025) Multi-omics reveals the inhibition of asexual reproduction in the jellyfish *Aurelia coerulea* polyp by *Bacillus pacificus*. Environmental Microbiome, 20, 78.

